# CryoSeek identification of glycofibrils with diverse compositions and structural assemblies

**DOI:** 10.1101/2025.09.30.679562

**Authors:** Zhangqiang Li, Tongtong Wang, Yitong Sun, Kui Xu, Wenze Huang, Qiangfeng Cliff Zhang, Chuangye Yan, Mingxu Hu, Nieng Yan

## Abstract

Last year, we reported CryoSeek, a research strategy that employs cryo-electron microscopy (cryo-EM) to discover novel bio-entities from any accessible source, supplemented with AI-facilitated data processing and bioinformatic analyses. Here we report CryoSeek characterization of five additional glycofibrils isolated from the <u>T</u>singhua <u>L</u>otus <u>P</u>ond (TLP), named TLP-IPT, TLP-12, TLP-3, TLP-2, and TLP-0, with overall resolutions ranging from 3.0-3.5 Å. These five glycofibrils, all covered with dense glycoshells, have decreasing ratios of the central protein components, with TLP-0 with no protein at all. IPT (immunoglobulin-like, plexins, transcription factors) refers to the tandem domain that constitutes the central filament of this type of glycofibrils. In TLP-12, the central cylindrical stem is made of a trimer of dodecapeptide repeats that weave to β-sheet ribbons. The central stem of TLP-3 is also a trimer, but of linear tripeptide repeats. TLP-2, similar to our previously reported TLP-4, only has a linear chain of dipeptide repeats, and glycosylation occurs to a phosphoserine in each repeat. Glycan-mediated interactions are essential for the assembly of all five glycofibrils. Our previous and present studies demonstrate the diversity in the high-order structure and folding of glycans.

## Introduction

Carbohydrates, or glycans, represent the most abundant organic molecules on Earth, and play essential roles in diverse biological processes (1–8). However, their structural characterization has long been exceptionally challenging. This difficulty mainly stems from the remarkable diversity and stereochemistry of monosaccharides, the complexity of branching patterns, and the absence of direct genetic templates encoding glycan biosynthesis (9–12). Consequently, high-resolution structures have primarily been determined for mono- or oligo-saccharides bound to proteins, or the several proximal sugar residues in the O- or N-linked glycans (13–18).

A serendipitous breakthrough in the determination of high-order structure of glycans occurred when we introduced an unconventional research strategy, CryoSeek, that employs cryo-EM to image minimally processed samples directly from natural sources or tissues (19–21). In the proof of principle experiments for CryoSeek, we collected and analyzed cryo-EM images for water specimen harvested from the Tsinghua Lotus Pond. The most prominent entities on the micrographs are fibrils of various diameters and curvatures. High-resolution 3D EM maps can be easily reconstructed for some of these fibrils that are made of repeating units. Automatic model building conveniently yielded pili-like structures. Sequences derived from the structures suggest that they are likely from uncharacterized bacteria (19).

However, the majority of the fibrils cannot be modeled through the auto-building methods (22, 23). Scrutiny of the map by experienced structural biologists confirmed that these fibrils all contain a large number of well-structured glycans whose model building is beyond the present capacity of CryoNet or ModelAngelos. Our initial attempts resulted in structural determination of two similar glycofibrils, which we name TLP-4a and TLP-4b. The nomenclature reflects both their origin from the <u>T</u>singhua <u>L</u>otus <u>P</u>ond and their central stems being a linear chain of tetrapeptide repeats. The glycoshell constitutes more than 90% of the molecular mass in these glycofibrils, and their structural assemblies are entirely supported by glycan mediated interactions (20, 21). The protein core is highly similar in TLP-4a and TLP-4b. Among the four residues in each repeat, two are conserved: a 3,4-dihydroxyproline (diHyp) bearing two O-linked glycans at 3-OH and 4-OH positions, and an adjacent serine/threonine (Ser/Thr) modified by a third O-linked glycan. The remaining two residues are less conserved. The principal differences between TLP-4a and TLP-4b lie in the number, linkages, and branching of sugar residues (20, 21).

Following upon our previous efforts, we hereby report discovery and structural determination of five additional types of TLP glycofibrils using CryoSeek. Whereas four of these fibrils feature protein cores densely decorated with glycans, a fifth one is composed of glycans only. In all cases, glycan-mediated interactions underlie fibril assembly. Our findings illustrate the diversity of glycofibrils and the structural roles of glycans, and further demonstrate the utility of CryoSeek for glycan structure characterization.

## Results

### CryoSeek discovery and structural overview of TLP glycofibrils

As a continuation of CryoSeek, procedures for sample preparation and cryo-EM data collection in this study were the same as previously described, and data processing and structural analysis were performed using similar protocols (19–21). In brief, fibrils with diameters larger than 5 nm and discernible two-dimensional (2D) features were classified into distinct groups for three-dimensional (3D) reconstruction (Fig. S1). Fibrils with diameters smaller than 5 nm were subjected to multiple rounds of 2D classification to enrich classes with well-defined features, from which one 3D reconstruction was obtained (Fig. S1). All the five resolved fibrils have overall resolutions of 3.0-3.5 Å (Fig. S2A). Many other fibrils also displayed distinctive 2D features, yet high-resolution 3D reconstructions could not be obtained, likely due to the limited unit numbers or structural heterogeneity (Fig. S2B).

Together with our previously reported TLP-1a/1b and TLP-4a/4b (19–21), we have now resolved nine distinct fibrils in the water samples collected from the Tsinghua Lotus Pond (Fig. 1A). Based on their molecular compositions, they fall into three classes: i) fibrils composed primarily of proteins; ii) fibrils containing both proteins and glycans, with glycans predominating in most cases; and iii) fibrils composed entirely of glycans. Following our previous nomenclature (19–21), we refer all fibrils with the prefix TLP. TLP-<u>1</u>a and TLP-<u>1</u>b were the <u>first</u> two structures resolved from the micrographs; the number in TLP-<u>4</u>a and <u>4</u>b refers to the basic unit of the <u>tetra</u>peptide in the central linear protein thread. In the present study, we name the five additional glycofibrils TLP-IPT, TLP-12, TLP-3, TLP-2, and TLP-0 based on their distinct protein components.

**Figure 1.**
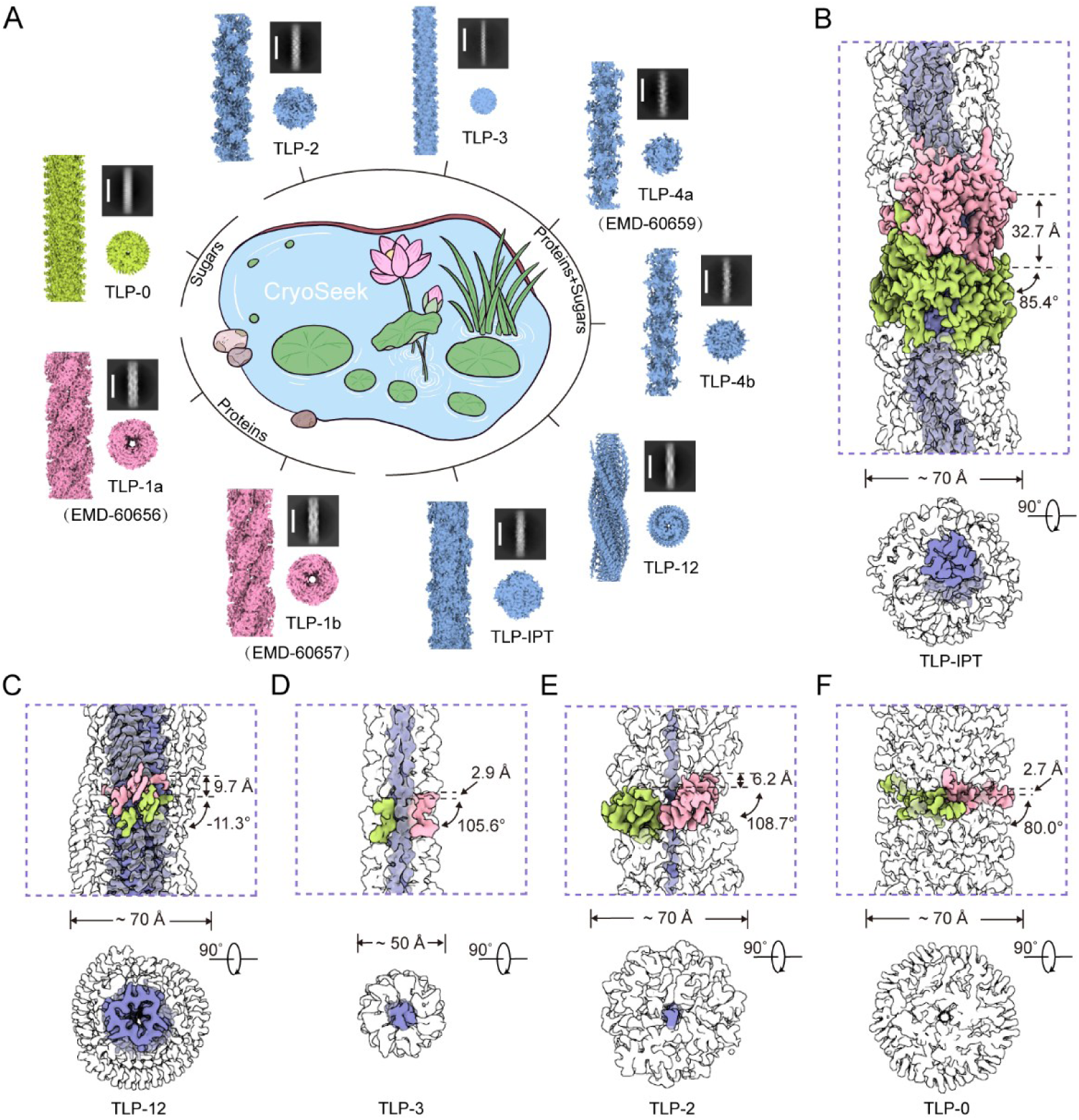
Discovery of various fibrils from the Tsinghua Lotus Pond via the CryoSeek strategy. (*A*) Cryo-EM overview of nine fibrils resolved at near-atomic resolution. Fibrils are classified into three groups based on their compositions: protein fibrils exemplified by TLP-1a and TLP-1b; glycofibrils with protein cores, including TLP-2/3/4a/4b/IPT; and a protein-free glycofibril, TLP-0. Two perpendicular views of each fibril are displayed together with their representative 2D class averages (scale bar 150 Å). TLP-1a/1b and TLP-4a/4b were reported previously, and the other five are illustrated in this study. (*B*-*F*) Cryo-EM density maps showing compositions and helical symmetries of TLP-IPT (*B*), TLP-12 (*C*), TLP-3 (*D*), TLP-2 (*E*) and TLP-0 (*F*). The densities for proteins and glycans are colored light purple and white, respectively; adjacent asymmetric units are highlighted green and pink. All EM maps are contoured at 6 σ throughout the manuscript if not otherwise indicated. All structural figures were prepared in ChimeraX (61).

TLP-IPT contains tandem immunoglobulin (Ig)-like domains that resemble beads on a string, except that the fibrillar assembly is fortified by glycans extending from adjacent protein domains. As will be discussed later, these Ig-like domains are, more specifically, the IPT (immunoglobulin-like, plexins, transcription factors) domains. Hence we call the glycofibril TLP-ITP. The distinguishable asymmetric unit of the TLP-IPT helical assembly gives to a helical rise of 32.7 Å and a helical twist of 85.4° (Fig. 1B). TLP-12 features a dodecapeptide repeat that forms a highly ordered triple β-sheet core, with a helical rise of 9.6 Å and a twist of –11.3° (Fig. 1C). TLP-3 and TLP-2 both contain linear peptide chains as their cores; TLP-3 is formed by triplex strands of tripeptide repeats, whereas TLP-2 consists of a single chain of dipeptide repeats, resembling the organization of TLP-4a/4b. The helical parameters are 2.9 Å, and 105.6° for TLP-3, and 6.2 Å, and 108.7° for TLP-2, respectively (Fig. 1D,E). TLP-0, which lacks a protein component and is composed solely of glycans, exhibits the helical parameters of 2.6 Å and –80.0° (Fig. 1F). Together, these structures underscore the compositional and organizational diversity of glycofibrils,.

### TLP-IPT is a glycofibril containing well-folded protein domain

Among the five glycofibrils described in this study, TLP-IPT contains the most easily discernible protein portion as the central axis. Automatic model building in CryoNet readily generated an atomic model for the protein domains (22), whereas the surrounding glycan densities had to be modeled manually (Fig. S3A). The automatically built model immediately revealed consecutive protein domains that each adopt an Ig-like fold, and specifically the IPT domain (Fig. 2A). Each domain, together with its attached glycans, constitutes one asymmetric unit. After manual inspection and correction of misfit regions, a refined protein core model of TLP-IPT was obtained (Fig. S3B, S3C). Local EM densities and the distances between glycosylated residues and attached sugars suggest all glycan chains in TLP-IPT may be O-linked to Ser/Thr. For simplicity, all glycosylated sites were provisionally assigned to Ser in the final model (Fig. 2A).

**Figure 2.**
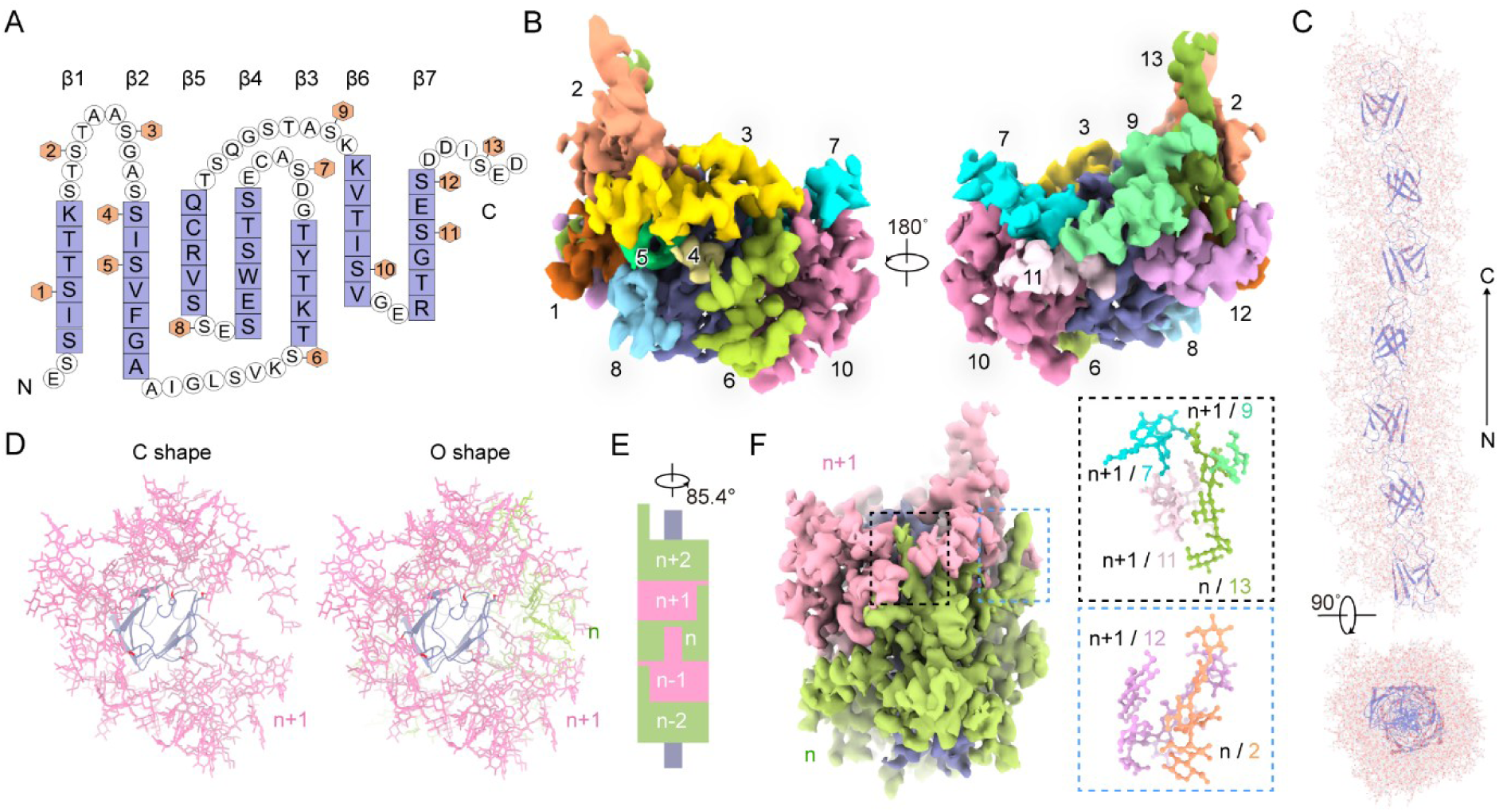
TLP-IPT comprises of consecutive IPT domains coated by dense glycans. (*A*) Topology of each IPT domain. Residues in light purple boxes represent β-strands and the glycans that are linked to Ser or Thr are indicated by orange hexagons. For simplicity, we assigned Ser for all the O-glycosylation sites. (*B*) The asymmetric unit of TLP-IPT consists of a single IPT domain modified with 13 glycan chains. The protein density is colored light purple, and each glycan chain is color-coded. (*C*) Structural model of a representative TLP-IPT segment that contains seven heavily glycosylated IPT domains. Protein repeats are colored light purple, and sugar residues are shown as light gray sticks with oxygen atoms colored red. (*D*) Resemblance of the “donor-strand exchange” assembly by the glycan chains in the adjacent repeats. Shown here are cross-section views. The glycan chains from repeat *n*+1 form a C-shaped structure, and the cleft is filled by the glycans from repeat n to complete a closed O ring. (*E*) Schematic illustration of the stacking of glycans that is reminiscent of the donor-strand exchange protein assembly observed in many bacterial pili. (*F*) Two glycan-mediated interfaces that complete the O ring formation. *Insets*: The first interface (upper) is formed by glycans 7, 9, and 11 from repeat *n+1* with glycan 13 from repeat *n*; the second interface (lower) is constituted by glycan 12 from repeat *n+1* and glycan 2 from repeat *n*. Glycans are colored the same as in panel *A*.

In each asymmetric unit, 13 glycan chains were identified. They attach to both loop regions and β-strands to form a dense glycoshell surrounding each protein domain (Fig. 2A,B). Although the current resolution was insufficient for accurate sugar residue assignment, we tentatively built structural models for these glycans to facilitate structural and functional assessment (Fig. 2C, Fig. S3C).

Extensive hydrogen-bonding and stacking interactions among sugar residues both within and between repeats of TLP-IPT play an important role in stabilizing the fibrillar structure. Notably, a unique interface was identified between repeat *n* and *n+1*. In the cross-sectional view, glycans linked to repeat *n+1* form a C-shaped contour. The cleft is filled by glycans from repeat *n* to complete an “O” ring (Fig. 2D). Such glycan-mediated inter-repeat interactions is reminiscent of the “donor-strand exchange” packing mechanism of many pili, in which such inter-repeat assembly is filled by protein segments though (19) (Fig. 2E). Two distinct hydrogen-bonding interfaces contribute to this organization: a major interface involving glycans 7, 9 and 11 from repeat *n+1*, and glycan 13 from repeat *n*, and a secondary interface formed by glycan 12 from repeat *n+1* and glycan 2 from repeat *n* (Fig. 2F).

IPT domains are typically found in cell surface receptors (24–26). Proteins such as HGFR and HST1R contain three to four IPT domains, and Plexin A1 contains up to six (25). Previous mass spectrometry characterizations also suggested the existence of numerous O-Man glycosylation sites on the IPT domains, although the detailed glycan compositions and branching patterns have remained unclear (27). While this information provides clues to the possible nature of TLP-IPT, the number of IPT domains in the resolved glycofibril, which is more than 600 nm in length, apparently exceeds 200, way more than those in the well-characterized receptors.

To investigate the potential origin and function of TLP-IPT, its structure-derived sequence was used to perform a BLAST search against the NCBI non-redundant (NR) database (28). A group of related proteins from *Baffinella frigidus*, a species that belongs to the class Cryptophyceae, stood out (Fig. S4A) (29). Among these, the sequence with GenBank ID KAJ1470089.1 was picked as a candidate for its highest BLAST score. Structural prediction of KAJ1470089.1 yielded a flexible filament composed of ten domains share a fold similar to the IPT domain (Fig. S4B). Sequence alignment of TLP-IPT with several repeat units of KAJ1470089.1 reveals high conservation at the glycosylation sites (Fig. S4C), and a TLP-IPT domain can be superimposed with one repeat of KAJ1470089.1 (residues 566-651) with the root-mean-square deviation of 0.7 Å over 86 Cα atoms (Fig. S4D). Nevertheless, the length of TLP-IPT indicates a sequence of at least 20,000 residues, which is much longer than the 1063 residues in KAJ1470089.1. Therefore, the host species, identity and function of TLP-IPT remain to be determined.

### TLP-12 is made of triple chains of heavily glycosylated dodecapeptide repeats

TLP-12 is also a fibril comprising a central protein stem coated with thick glycans (Fig1A,C). Structural analysis reveals that the stem of TLP-12 is formed by three polypeptide chains that intertwine and spiral upward in parallel, forming a central cylinder with a triangular cross-section. On each side of the triangle are two bulky densities corresponding to glycans (Fig. 3A,B). The larger and smaller glycans constitute two parallel helical ridges that wrap around the fibril. The uncoated protein region, which is quite narrow, appears as a groove in the EM map (Fig. 3C).

**Figure 3.**
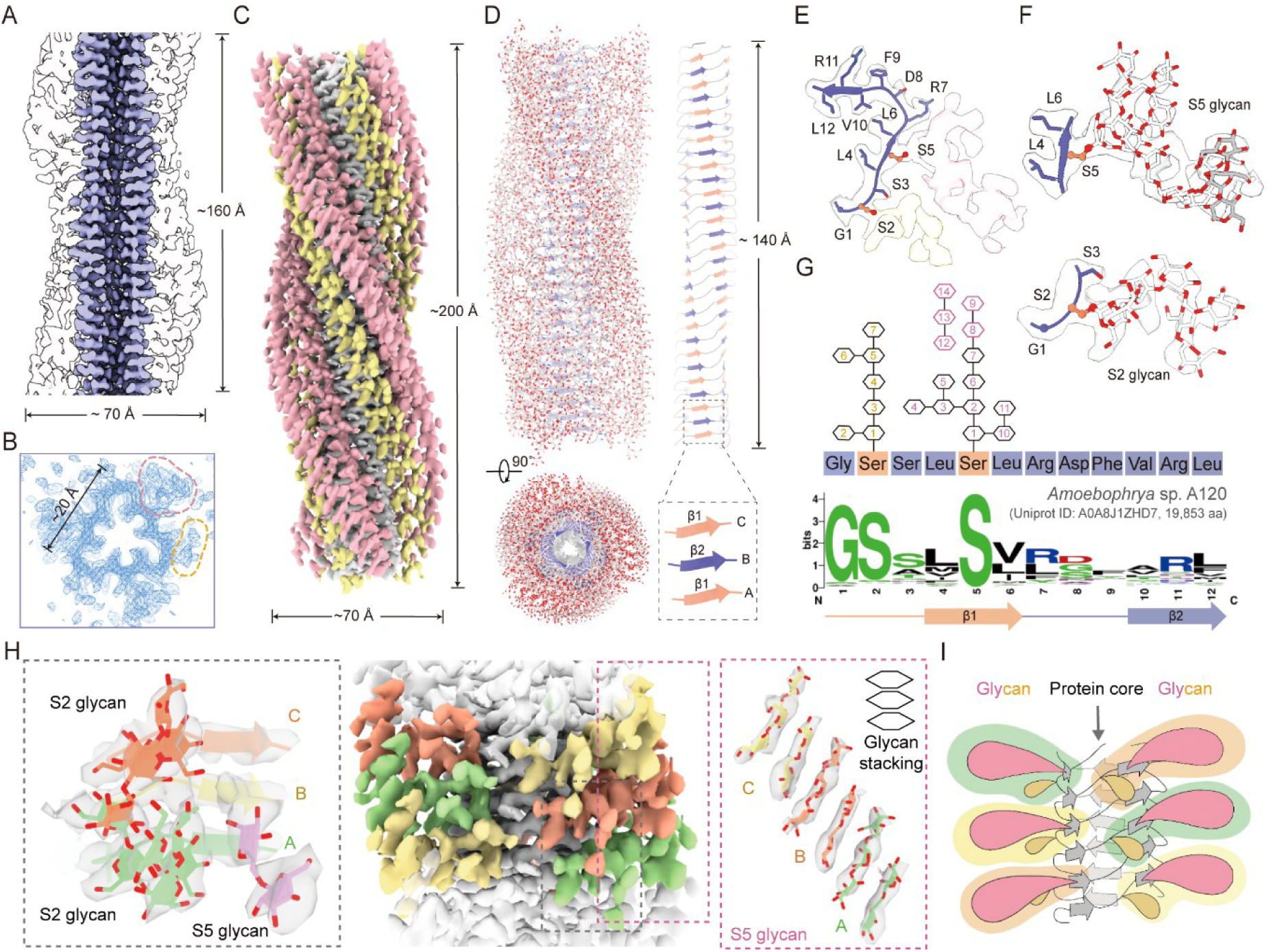
The TLP-12 fibril is a trimer of heavily glycosylated dodecapeptide repeats. (*A*) TLP-12 fibril comprises central protein repeats coated with thick glycan shell. The map is contoured at 6 σ. The central protein portion is colored light purple, and the surrounding white densities belong to glycans. (*B*) C3 symmetry of TLP-12. Shown here is a cross-section of TLP-12. Two glycosylation densities, one bigger and one smaller indicated respectively with red and orange dashed lines, are well resolved on each side of the triangle. The density is contoured at 7.5 σ in COOT. (*C*) Glycans define the ridge and groove along the spiral fibril. The larger and smaller glycans attaching to each repeat are colored pink and yellow, respectively. (*D*) Trimeric structure of a segment of the TLP-12. *Left*: Two perpendicular views of the structure of a TLP-12 segment. The protein repeats are colored light purple, and the sugar moieties are shown as gray sticks with oxygen atoms colored red. *Right*: The central protein shaft is made of interwoven trimer. The two β strands in each dodecapeptide repeat are colored orange and light purple in all three monomers. An enlarged view of the parallel β strands from the three monomers A/B/C are shown in the inset. (*E*) Model building of a representative dodecapeptide repeat. The smaller and larger glycans are linked to the 2^nd^ and 5^th^ positions, respectively. Ser is tentatively assigned to both positions, but we cannot exclude the possibility of Thr at either or both positions. (*F*) Tentative assignment of glycosyl moieties, named S2 glycan and S5 glycan, into the density. The linkage of three connected sugar moieties, colored gray, cannot be identified, but their position suggests that they most likely belong to the S5 glycan. Sugar types and configurations cannot be precisely determined due to the resolution limitation of the EM map and the lack of additional evidence. (*G*) Conservation analysis of the dodecapeptide repeats of a protein from a species of dinoflagellate, *Amoebophrya* sp. A120. Shown below is a representative protein with Uniprot ID of A0A8J1ZHD7, which is predicted to be 19,853-residue long, with 1,263 GSxxSx_7_ repeats. The degenerate sequence above the motif logo is consistent with the density shown in panel E. The sugar moieties linked to the 2^nd^ and 5^th^ positions are represented by hexagons to indicate the ambiguity in their identity. (*H*) Glycan interactions in TLP-12. The glycans in strands A/B/C are colored green, light blue, and orange, respectively. *Left inset*: S2 glycan from strand B may form extensive hydrogen bonds with S5 glycan from strand B and S2 glycan from strand A. *Right inset*: Glycan stacking of S5 glycans at the edge of the triangle of TLP-12. (*I*) A schematic cartoon to illustrate the structural assembly of TLP-12. Apart from the co-folded trimeric protein stem, extensive contacts are mediated by glycans.

In contrast to TLP-IPT, automatic model building for TLP-12 was unsuccessful despite its proteinaceous core. To address this, glycan densities were manually erased before importing the remaining densities into CryoNet (22). The resulting model revealed a unique repeating dodecapeptide structural motif in each protein chain, which provide the basis for the designation of TLP-12 (Fig. 3D, Fig. S5A). Each dodecapeptide repeat spans two sides of a triangle, with a short β-strand on each side (β1, residues 4-6; β2, residues 10-12). Parallel β-strands from the three protein chains, following the repetitive pattern A/β1-B/β2-C/β1-A/β2-B/β1-C/β2, weave an elongated and narrow β-sheet ribbon that spirals around the central axis. The sugar moieties were then modeled manually according to the local densities (Fig. 3D).

An invariant Gly was defined as the first residue of the repeat. The smaller and larger glycans are linked to the residues at the 2^nd^ and 5^th^ positions, respectively (Fig. 3E,F). In most repeats, both positions are occupied by Ser. We hereafter refer to them as S2 glycan and S5 glycan. The averaged resolution of 3.0 Å does not support accurate assignment of the glycosyls. We tentatively modeled 7 and 11 sugar moieties for the S2 and S5 glycans, respectively. Additionally, a stretch of density that can be fitted with three glycosyls appears to be part of the S5 glycan, although the linkage is invisible (Fig. 3F).

To explore the potential origins of TLP-12, the bseek tool in CryoNet (https://cryonet.ai/bseek) was applied to search for experimental and predicted structures in PDB, AFDB, and the Evolutionary Scale Modeling Metagenomic Atlas (ESM30) that match the auto-modeled TLP-12 backbone (22, 30–33). A structural model from ESM30 (MGnify ID: MGYP001210373462) exhibited a similar fold to a single TLP-12 chain (Fig. S5B,C,D). This sequence was used to BLAST for additional homologues in the NCBI NR database, identifying a group of proteins, all from the parasitic dinoflagellate *Amoebophrya* (34, 35). Most are large proteins, consisting approximately 20,000 residues. For instance, a protein with Uniprot ID A0A8J1ZHD7 from *Amoebophrya* sp. A120 contains 19,853 residues.

The dodecapeptide repeats featuring G1, S2, and S5 are highly conserved across these candidates, and the conserved residues of A0A8J1ZHD7 align well with the averaged density map (Fig. 3E,G) (36). Aided by these clues, we manually refined the auto-built model. In the final structure of a TLP-12 segment spanning 14 nm, 648 residues fold into 18 repeats in each monomeric chain, and are modified with 1,134 sugar moieties (Fig. 3D).

A *FIMO* (*Find Individual Motif*) search using the dodecapeptide repeat further identified ∼ 150 protein candidates from 6 species or strains within the Alveolate clade, although none of them has characterized functions (Fig. S5E) (37, 38). Except for the free-living *Polarella glacialis*, the other species are either parasitic or symbiotic (39–42).

To further probe the structural and functional implications of these candidates, we selected A0A8J1ZHD7 as a representative sequence for InterPro analysis. Domain prediction revealed numerous concentrated dodecapeptide repeats in A0A8J1ZHD7 and indicated the presence of an ICA (intramolecular chaperone autocleavage) domain at its C terminus (Fig. S5F,G). The ICA domain has been well studied in mammals and phages, and structural analyses show that it promotes the folding of three β-strands into a triple β-helix (43–45). Comparing the predicted ICA structures from TLP-12 candidates with deposited ICA domains exhibited high structural similarity (Fig. S5H), suggesting that the C terminus is essential in mediating triple β-helix folding in TLP-12 (Fig. S5F,G).

In addition to the triple β-helix, the decorated glycans are also important for TLP-12 assembly (Fig. 3H,I). The S2 and S5 glycans within the same repeat have a loose contact through hydrogen bonds between two adjacent sugar moieties, one from each glycan (Fig. 3E). Furthermore, S2 glycans from alternating monomers, like A and C or B and A, are clustered through polar interactions (Fig. 3H, left). A stunning pattern is the stacking of di-glycosyls from S5 glycans in adjacent monomers (Fig. 3H, right). The stacked di-glycosyls may contribute to the stabilization of the glycocalyx assembly, hence explaining the better resolution of the tri-glycosyls, assigned sequential numbers 12-14, although their covalent linkage to the rest of the glycan is invisible in the EM map (Fig. 3G).

### Glycofibrils with linear peptide cores

Among the fibrils identified, TLP-3 and TLP-2, together with our previously reported TLP-4a/4b (20, 21), represent a distinct class of glycofibrils whose protein cores comprise linear peptides with short repetitive sequences. In contrast to TLP-IPT and TLP-12, whose folded protein domains enabled automatic model building, these glycofibrils with linear peptide cores could not be modeled directly and required manual interpretation of the cryo-EM densities.

TLP-3 is a trimeric fibril with the diameter of approximately 4 nm (Fig. 1A,D). Its protein core is made of triplex spiral that is completely covered by thick glycans (Fig. 4A). Each protein strand comprises of repeating tripeptide units, in which the first two are variable residues with small side chains and the third corresponds to O-glycosylated Ser/Thr. For simplicity, we modeled the repeats as Gly-Gly-Ser (Fig. 4B,C). Eight sugar residues were modeled into the EM densities in each repeat (Fig. 4C,D). The sugar residue that is directly linked to Ser (sugar 1) has three additional branches: a single sugar 5 sandwiched by sugars 2-4 in branch 1 and sugars 6-8 constitute branch 2.

**Figure 4.**
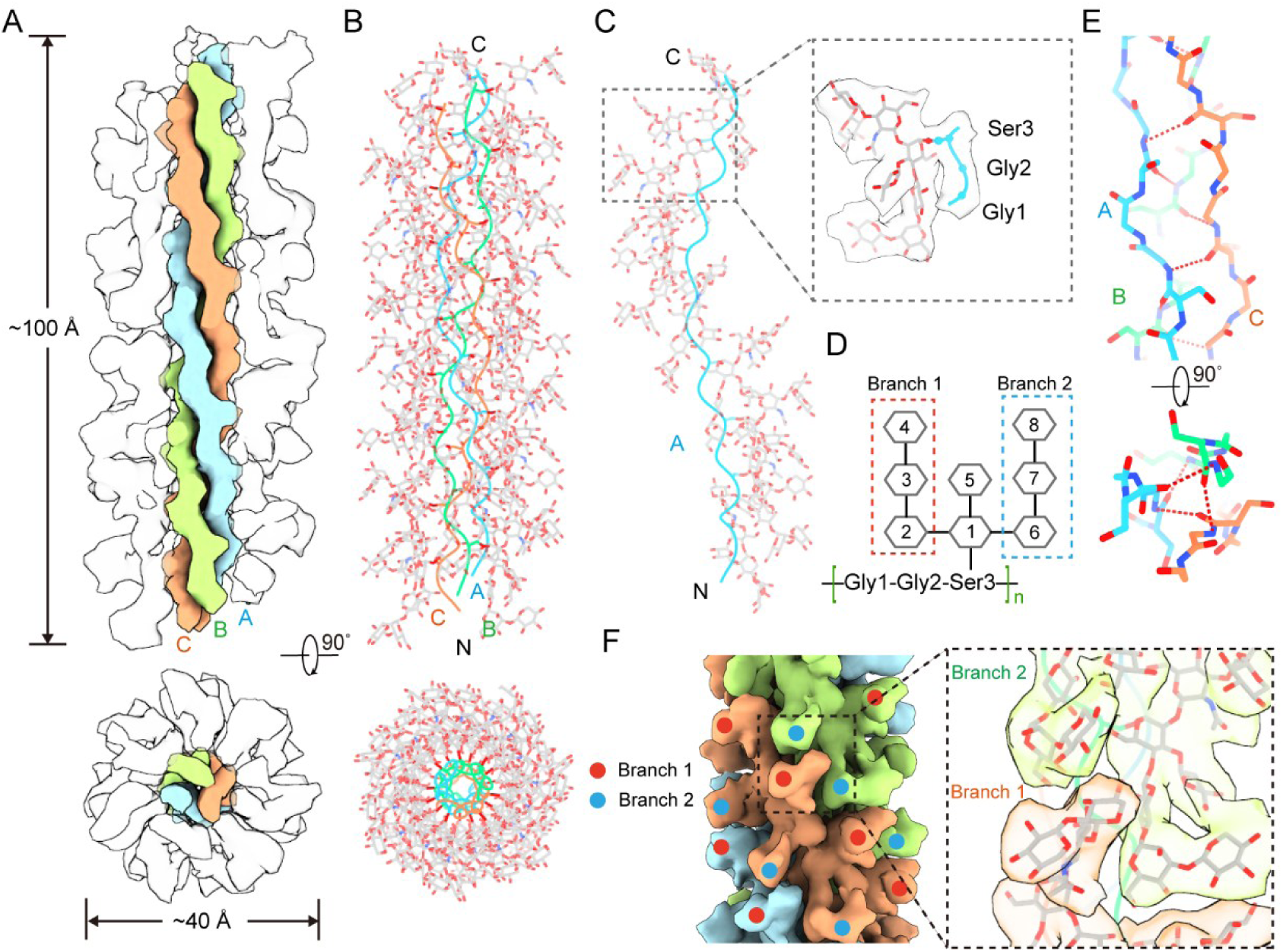
TLP-3 is a triplex of linear tripeptide repeats coated with glycoshell. (*A*) Trimeric structure of TLP-3. Shown here are two perpendicular views of the 3D reconstruction of TLP-3. The triplex strands of the tripeptide repeats are colored blue, green, and orange, respectively. Surrounding glycan densities are colored white. (*B*) Structural model of a representative TLP-3 segment. (*C*) A single strand of TLP-3. *Inset*: Each repeat consists of three residues, tentatively assigned with Gly-Gly-Ser. Eight sugar residues were fitted into the glycan density that is linked to Ser. (*D*) Schematic illustration of a single repeat. Sugar moieties linked to the Ser are represented by uncolored hexagons to indicate ambiguity in their identity. (*E*) Inter-strand interactions mediated by hydrogen bonds between backbone groups in TLP-3. A hydrogen-bond network is formed between the carboxyl group of Ser3 in one chain and the amide group of Gly1 in the neighboring chain. (*F*) Glycan-mediated interactions in TLP-3. Glycans from adjacent repeats interlock in a helical fashion to form the glycoshell. Packing details are shown in the inset.

Helical stabilization arises from both protein- and glycan-mediated interactions. Hydrogen bonds between the carboxyl group of Ser3 in one chain and the amide group of Gly1 in the neighboring chain stabilize the protein core, while glycans from the adjacent repeats interlock to shape the surrounding glycan coat (Fig. 4E,F).

For TLP-2, the helical rise (6.2 Å) is half of the previously reported TLP-4a/4b (12.4 Å), indicating a repeat unit of dipeptide (Fig. 1A,E). Manual checking and modeling confirmed this analysis, each repeat having one conserved glycosylated residue and one variable residue. The cryo-EM density did not support conventional N- or O-glycosylation linkages. Although atypical O-glycosylation on hydroxylysine satisfied the length requirement for glycosidic bond, it did not fit into the local density well. Instead, this density was consistent with phosphoglycosylation, such as Ser-linked GlcNAc-1-PO4 described in *Dictyostelium discoideum* Proteinase I (Fig. 5A) (46, 47). We therefore assigned the repeat as Ala-Ser for better illustration. In the final model, each Ser is linked via phosphodiester bonds to an extended glycan chain (Fig. 5B,C). Glycan-mediated assembly of TLP-2 closely resembles that of TLP-4a/4b. Following the helical twist, the asymmetric repeats of TLP-2 are indexed as *n*, *n+1*, *n+2*, and *n+3*. Under this scheme, inter-repeat interactions are observed between *n* and *n+1*, and between *n* and *n+3* (Fig. 5D). Complementary packing and hydrogen-bonding networks among adjacent glycans collectively stabilize the helical structure of TLP-2.

**Figure 5.**
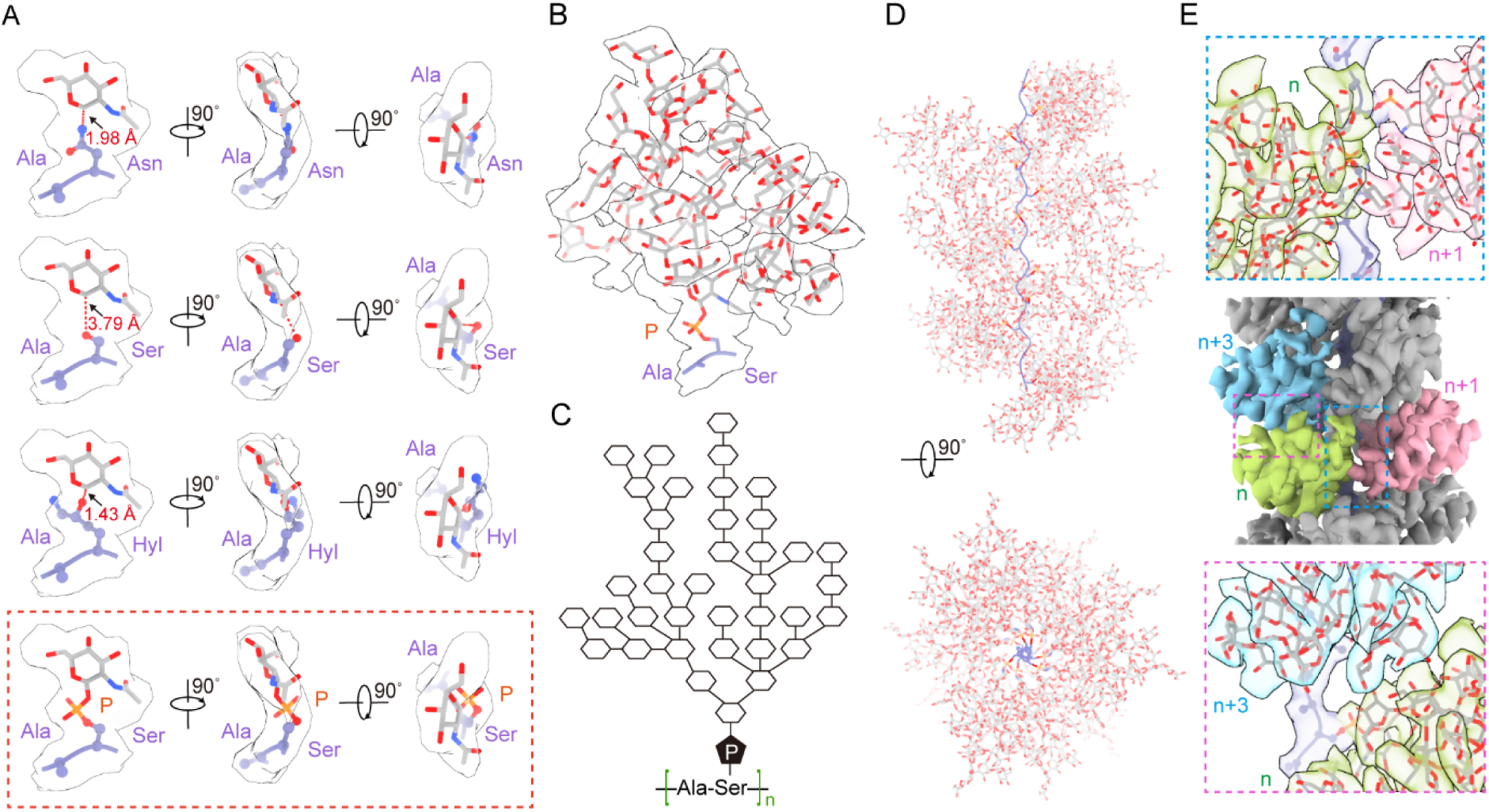
TLP-2 is a glycan-coated linear chain of dipeptide repeats. (*A*) Sugar chain linked to a phosphoserine fits the density best among different options. Show here are our attempts to model in different glycosylation types into the cryo-EM density that is contiguous to the protein moiety. Ser-linked GlcNAc-1-PO_4_, instead of N-linked Asn or hydroxylysine, best matches the local density. (*B*) Atomic model of the asymmetric unit of TLP-2. Ala and Ser were tentatively assigned to each repeat, and 38 generic sugar residues were modeled to the glycan linked to phosphoserine. (*C*) Schematic illustration of glycans linked to an asymmetric. Sugar moieties are represented by uncolored hexagons to indicate the ambiguity in their identity. (*D*) Structure of a TLP-2 segment shown in two perpendicular views. (*E*) Structural role of glycan-mediated interactions in the assembly of TLP-2. Hydrogen bonds formed by the glycans from repeat *n* and *n*+1, and from repeat *n* and *n*+3, define the assembly of TLP-2.

TLP-3 and TLP-2, as well as the previously reported TLP-4a/4b, highlight the diversity of glycofibrils with linear peptide cores. All of these fibrils have short peptide repeats as minimal scaffolds, and rely on extensive glycosylation to stabilize their high-order assembly. Although sequence BLAST for TLP-4a/4b yielded important clues to their origins, a similar search for TLP-3 and TLP-2 failed to retrieve useful hits, possibly due to limited sequence deposition as well as the inherent challenges in sequencing the large number of tripeptide and dipeptide repeats.

### TLP-0 is composed exclusively of glycans

Unlike the other resolved TLP fibrils, the 3D reconstruction of the last fibril revealed no densities corresponding to amino acid residues. Accordingly, we designated this fibril as TLP-0, with “0” indicating the absence of protein. As the chirality and orientational information of TLP-0 is completely intractable, we modelled common stereochemically correct monosaccharides into densities of TLP-0, (Fig. 6A). The core of TLP-0 consists of a spiral tri-glycosyl repeat (Fig. 6A). With a local resolution of nearly 2.8 Å in the core region, we modeled the tri-glycosyl repeat with high confidence. Fucα1-3GalNAcα1-3Man fits the EM density well, with Man of repeat *n* linked to Fuc of repeat *n-1* through an α1-3 linkage (Fig. 6B).

**Figure 6.**
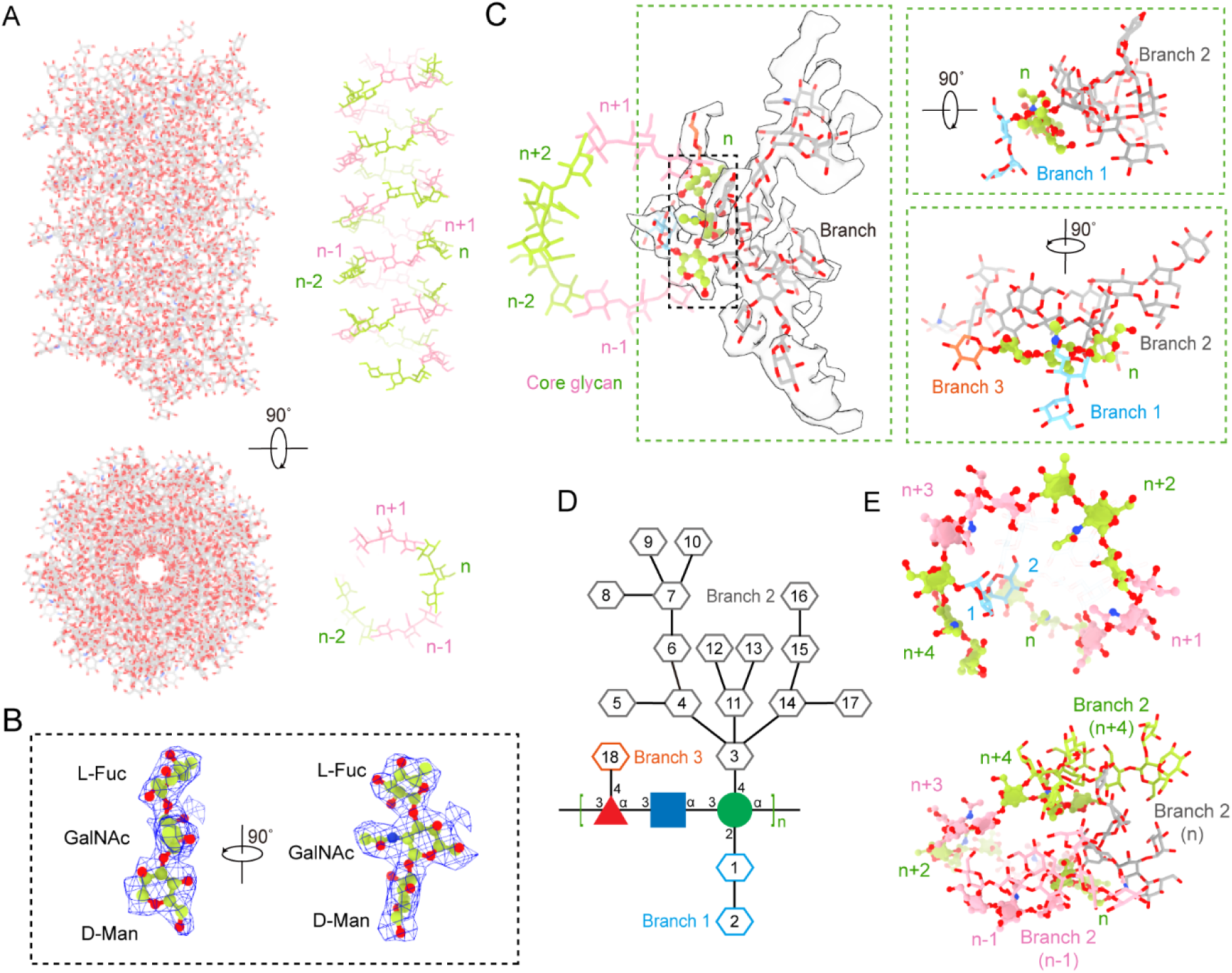
TLP-0 is formed entirely by glycans. (*A*) TLP-0 is a protein-free fibril with a core of tri-glycosyl repeats. Two perpendicular views of a segment of tri-glycosyl repeats in the core are shown on the right. For clarity, the tri-glycosyl repeats are alternately colored green and pink. (*B*) Local densities for the core tri-glycosyl repeat. L-Fuc, GalNac, and D-Man fit the densities well. (*C*) Structural organization of TLP-0. In one asymmetric unit, three branches are linked to the tri-glycosyl-repeat core. Branch 1, facing the interior of TLP-0, is linked to Man. Branch 2 and Branch 3, facing the exterior of TLP-0, are linked to Man and Fuc, respectively. The three branches are colored blue, gray, and orange, respectively. (*D*) Schematic illustration of an asymmetric unit of TLP-0. The core of TLP-0 is Fucα1-3GalNAcα1-3Man with high confidence, whereas the other 18 sugar moieties are presented by hexagons to indicate ambiguity in their identity. The red triangle, blue rectangle, and the green circle stand for Fuc, GlcNAc, and Man, respectively, according to monosaccharide symbol nomenclature. (*E*) Two critical interfaces responsible for the assembly of TLP-0. Glycan 2 from repeat *n+4* interacts with Man in repeat *n* via glycan stacking. Multiple hydrogen bonds form between Banch 2 in repeat *n* and the corresponding branches in repeat *n-1* and *n+4*. These glycan-mediated interactions determine the structural assembly of TLP-0.

There are three glycan branches attached to each of the core tri-glycosyl repeat, with branch 1 facing inward and branches 2 and 3 facing outward (Fig. 6C). Their relative positions and branching patterns were tentatively modeled. In total, 18 sugars were built onto the tri-glycosyl repeat: 2 sugars in branch 1 linked to Man via an α1-2 linkage, 15 sugars in branch 2 linked to Man via an α1-4 linkage, and 1 sugar in branch 3 linked to Fuc via an α1-4 linkage. Together, these glycans define one asymmetric unit of TLP-0 (Fig. 6C,D).

To better describe the interactions among sugar residues, we indexed the asymmetric units along the spiral direction. On the inside of TLP-0, sugar 2 of branch 1 in repeat *n+3* stacks against Man of repeat *n*. On the outside, branch 2 of repeat *n* engages in multiple interactions with branch 2 of repeats *n-1* and *n+4* (Fig. 6E).

In the absence of protein sequence, it is impossible to retrieve the origin of TLP-0 using bioinformatic analysis. The all-sugar composition of TLP-0 reminds us of starch, cellulose and glycogen (48, 49). Starch and glycogen serve as energy storage substances in many organisms (50, 51), and cellulose is the major structural component of cell walls in plants and algae (52). All three consists solely of glucose, differing only in linkage and branching patterns. By contrast, TLP-0 has a more complex composition, with its core built from three kinds of sugars. Discovery of this all-sugar fibril immediately raises many questions that await comprehensive investigations.

## Discussion

In this study, we report the CryoSeek identification of five additional glycofibrils from the water samples collected in the Tsinghua Lotus Pond. It should be noted that the sugar residues modeled into the peripheral densities may not reflect their real identities due to the moderate resolution. The decreased resolutions on periphery suggest a higher glycan-to-protein mass ratio in these glycofibrils than that reflected in the structural models (Fig. S6).

Although their physiological functions remain unclear, these glycofibrils may in general provide a strategy for carbon storage outside the cell body (48, 49). Unlike starch and glycogen, which accumulate intracellularly, or cellulose, which defines the cell wall, the glycofibrils reported here and previously extend into the extracellular milieu, offering potentially unlimited space for carbon storage. Such fibrils could act as nutrient reservoirs for the producing organisms or be utilized by other species. In addition, their distinct sugar-to-protein ratios may help regulate carbon-nitrogen balance. Considering the links of TLP-IPT to Cryptomonads (53), and TLP-12 to Alveolate (40), it is also possible that some glycofibrils function in host-parasite interactions or mediate survival strategies in specific ecological niches.

Our findings highlight the unique role of glycan-mediated interactions in determining high-order structural assembly. In contrast to the conventional paradigm in which proteins determine high-order architecture and glycans play minor roles, the glycofibrils described here, particularly TLP-0, demonstrate that glycans can also dominate structural folding.

Several technical challenges and limitations remain in our present study. For fibrils with well-folded protein domains (e.g. TLP-1a/1b and TLP-IPT), the handedness of the 3D reconstructions can be validated based on the structure of the protein folds. In contrast, for glycofibrils with linear peptide repeats or even without protein, handedness cannot be resolved empirically. In an accompanying study, we report the method for determining the handedness of glycofibrils to address this issue (54, 55).

In sum, our results, together with previous analyses of TLP-1 and TLP-4, demonstrate the utility of CryoSeek for discovering and characterizing novel bio-entities. Our studies not only expand the repertoire of known fibrous assemblies, but also raise fundamental questions about the structural roles of glycans, their evolution, biogenesis, and physiological functions. They exemplify how unexpected molecular architectures can transform our understanding of carbohydrates in biology. In a sense, the CryoSeek strategy may represent “forward structural biology”, a structure-first paradigm for biological discovery.

## Materials and Methods

### Data processing

Methods regarding sample preparation and data collection have been previously described (19, 20). A total of 18,037 movies were collected and subjected to motion correction using Motion Cor2, followed by patch CTF estimation in cryoSPARC (56). Particles were picked with the filament tracer tool. After inspecting particle picks, 8,930,454 particles were extracted with a rescaled box size of 80 pixels from the original 320 pixels (bin4). After several rounds of 2D classifications, particle sets corresponding to the five fibrils were identified: 248,923 particles for TLP-IPT, 96,204 particles for TLP-12, 32,488 particles for TLP-2 and 37,889 particles for TLP-0. Fibrils with diameters greater than 5 nm and smaller than 5 nm were separately classified at 2D level, as features for smaller fibrils were harder to stand out.

For all fibrils, ab-initio reconstruction was first carried out. The resulting volumes were then subjected sequentially to helical refinement without predefined helical symmetries, symmetry search, helical refinement with calculated helical symmetries, followed by heterogenous refinement to further enrich good particles. Before the final refinement, EM densities were manually inspected to ensure that the helical parameters between asymmetric units were consistent with the initial calculations. When discrepancies were observed, an additional round of symmetry search with restricted ranges would be performed and the optimized parameters would be imposed for the final helical refinement.

Among the five fibrils, TLP-IPT required re-extraction at 384 pixels (bin1) due to its large asymmetric unit. Helical refinement and symmetry search yielded a rise (Δ*z*) of 32.7 Å and a twist (Δ *Φ*) of 85.1°, with 70,391 particles contributing to the final 3.3 Å map. For TLP-12, ab-initio reconstruction at bin4 failed, so bin1 particles were used. Helical refinement and symmetry search first suggested parameters of 19.4 Å and -23.3°, but manual inspection showed the asymmetric unit was half this size, and the parameter optimization converged on 9.7 Å and -11.3°. A final set of 43,778 particles produced a 3.0 Å map. The initial model of TLP-3 was processed at 256 pixels (bin1). Preliminary parameters were defined as 5.9 Å and -148.5° and later refined to 2.9 Å and 105.6°, with 28,629 particles yielding a 3.5 Å map. TLP-2 underwent ab-initio reconstruction at bin4 (80 pixels from 320 pixels), followed by helical refinement at bin1 (320 pixels). The initial parameters were 12.5 Å and 145.7° and optimized to 6.2 Å and 108.7°. 18,637 particles were preserved to generate the 3.5 Å map. For TLP-0, bin4 (320 pixels) ab-initio reconstruction followed by bin1 (320 pixels) re-extraction for helical refinement gave preliminary parameters of 13.4 Å and 40.0°, which were refined to 2.7 Å and 80.0°. With 35,270 particles retained, the final reconstruction reached 3.1 Å.

For TLP-IPT and TLP-12, handedness was determined based on their respective protein core. Regarding TLP-2, the handedness of the related fibril, which was derived from a distinct natural freshwater source, was determined using a novel algorithm (Ahaha) in our companion study (54). Accordingly, we adopted those helical parameters as a reference for TLP-2 in this work. In contrast, The handedness of TLP-3 and TLP-0 remains undetermined.

The reported resolutions for the five additional fibrils in this study were calculated based on the gold-standard Fourier shell correlation (FSC) 0.143 criterion using cryoSPARC (56, 57).

### Model building and refinement

Model building of TLP-IPT/12/3/2/0 was carried out based on their corresponding reconstruction maps. The protein portions of TLP-IPT and TLP-12 were auto-built with CryoNet and then manually corrected. Since the protein portions of TLP-3 and TLP-2 are simple, we built them manually based on the EM densities. As the precise identities of the glycans could not be determined, we constructed hexoses or pentoses according to the EM maps. Finally, the atomic models of TLP-IPT/12/3/2/0 were manually built in COOT (58).

The final models of TLP-IPT/12/3/2/0 were refined using PHENIX with secondary structure and geometry restraints in real space (59). The structures were validated through examination of the Clash scores, MolProbity scores, and statistics of the Ramachandran plots in PHENIX (59, 60).

## Author contributions

N.Y. and Z.L. conceived the project. Z.L., T.W. and Y.S. performed experiments and collected cryo-EM data. Z.L., T.W., Y.S. and M.H. processed the cryo-EM data and determined the structure. Z.L., T.W., Y.S., K.X., M.H. and N.Y. did structural analysis. T.W. and W.H. conducted bioinformatic analysis. N.Y., Z.L. and T.W. wrote the paper.

## Acknowledgements

We thank Dr. Xiaomin Li and Dr. Jianlin Lei for technical support during EM image acquisition. We thank the Tsinghua University Branch of China National Center for Protein Sciences (Beijing) for providing the proteomics facility and cryo-EM facility support. We thank the computational facility support on the cluster of Bio-Computing Platform (Tsinghua University Branch of China National Center for Protein Sciences Beijing). This work was funded by the National Natural Science Foundation of China (project 92478205 and 32330052 to N.Y.).

## References

1. A. Varki, Biological roles of glycans. Glycobiology 27, 3–49 (2017).

2. A. Varki, P. Gagneux, “Biological Functions of Glycans” in Essentials of Glycobiology, A. Varki et al., Eds. (Cold Spring Harbor (NY), 2015), 10.1101/glycobiology.3e.007, pp. 77–88.

3. A. Varki, Biological roles of oligosaccharides: all of the theories are correct. Glycobiology 3, 97–130 (1993).

4. D. Cosgrove, Growth of the plant cell wall. Nat Rev Mol Cell Biol 6, 850–861 (2005).

5. Y. Amazaki, H. Nguyen, R. Okamoto, Y. Maki, Y. Kajihara, Effects of N-Glycans on Glycoprotein Folding and Protein Dynamics. Adv Exp Med Biol 1104, 1–19 (2018).

6. G. Rabinovich, Y. van Kooyk, B. Cobb, Glycobiology of immune responses. Ann N Y Acad Sci 1253, 1–15 (2012).

7. R. Flynn et al., Small RNAs are modified with N-glycans and displayed on the surface of living cells. Cell 184, 3109–3124 e3122 (2021).

8. J. Huang et al., Structure-guided discovery of protein and glycan components in native mastigonemes. Cell 187, 1733–1744 e1712 (2024).

9. R. Cummings, J. Pierce, The challenge and promise of glycomics. Chem Biol 21, 1–15 (2014).

10. S. Tommasone et al., The challenges of glycan recognition with natural and artificial receptors. Chem Soc Rev 48, 5488–5505 (2019).

11. H. Wilkinson, R. Saldova, Current Methods for the Characterization of O-Glycans. J Proteome Res 19, 3890–3905 (2020).

12. P. de Haas, W. Hendriks, D. Lefeber, A. Cambi, Biological and Technical Challenges in Unraveling the Role of N-Glycans in Immune Receptor Regulation. Front Chem 8, 55 (2020).

13. D. Deng et al., Crystal structure of the human glucose transporter GLUT1. Nature 510, 121–125 (2014).

14. D. Deng et al., Molecular basis of ligand recognition and transport by glucose transporters. Nature 526, 391–396 (2015).

15. T. Xie et al., Crystal structure of the γ-secretase component nicastrin. Proc Natl Acad Sci U S A 111, 13349–13354 (2014).

16. A. Walls et al., Glycan shield and epitope masking of a coronavirus spike protein observed by cryo-electron microscopy. Nat Struct Mol Biol 23, 899–905 (2016).

17. W. Williams et al., Fab-dimerized glycan-reactive antibodies are a structural category of natural antibodies. Cell 184, 2955–2972 e2925 (2021).

18. X. Xiong et al., Glycan Shield and Fusion Activation of a Deltacoronavirus Spike Glycoprotein Fine-Tuned for Enteric Infections. J Virol 92 (2018).

19. T. Wang et al., CryoSeek: A strategy for bioentity discovery using cryoelectron microscopy. Proc Natl Acad Sci U S A 121, e2417046121 (2024).

20. T. Wang et al., CryoSeek II: Cryo-EM analysis of glycofibrils from freshwater reveals well-structured glycans coating linear tetrapeptide repeats. Proc Natl Acad Sci U S A in press (2024).

21. T. Wang, Y. Sun, Z. Li, N. Yan, The 8-nm spaghetti: well-structured glycans coating linear tetrapeptide repeats discovered from freshwater with CryoSeek. bioRxiv, 2024.2012. 2015.627649 (2024).

22. K. Xu, Z. Wang, J. Shi, H. Li, Q. Zhang (2019) A2-net: Molecular structure estimation from cryo-em density volumes. in Proceedings of the AAAI Conference on Artificial Intelligence, pp 1230–1237.

23. K. Jamali et al., Automated model building and protein identification in cryo-EM maps. Nature 628, 450–457 (2024).

24. A. Williams, A. Barclay, The immunoglobulin superfamily--domains for cell surface recognition. Annu Rev Immunol 6, 381–405 (1988).

25. P. Bork, T. Doerks, T. Springer, B. Snel, Domains in plexins: links to integrins and transcription factors. Trends Biochem Sci 24, 261–263 (1999).

26. Q. Ma, K. Zhang, S. Guin, Y. Zhou, M. Wang, Deletion or insertion in the first immunoglobulin-plexin-transcription (IPT) domain differentially regulates expression and tumorigenic activities of RON receptor Tyrosine Kinase. Mol Cancer 9, 307 (2010).

27. M. Vester-Christensen et al., Mining the O-mannose glycoproteome reveals cadherins as major O-mannosylated glycoproteins. Proc Natl Acad Sci U S A 110, 21018–21023 (2013).

28. J. Ye, S. McGinnis, T. Madden, BLAST: improvements for better sequence analysis. Nucleic Acids Res 34, W6–9 (2006).

29. N. Daugbjerg, A. Norlin, C. Lovejoy, Baffinella frigidus gen. et sp. nov. (Baffinellaceae fam. nov., Cryptophyceae) from Baffin Bay: Morphology, pigment profile, phylogeny, and growth rate response to three abiotic factors. J Phycol 54, 665–680 (2018).

30. Z. Guo, Y. Wang, G. Ou, Utilizing the scale-invariant feature transform algorithm to align distance matrices facilitates systematic protein structure comparison. Bioinformatics 40 (2024).

31. F. Bernstein et al., The Protein Data Bank: a computer-based archival file for macromolecular structures. J Mol Biol 112, 535–542 (1977).

32. J. Jumper et al., Highly accurate protein structure prediction with AlphaFold. Nature 596, 583–589 (2021).

33. Z. Lin et al., Evolutionary-scale prediction of atomic-level protein structure with a language model. Science 379, 1123–1130 (2023).

34. L. Guillou, J. Szymczak, C. Alves-de-Souza, Amoebophrya ceratii. Trends Parasitol 39, 152–153 (2023).

35. S. Kim et al., Genetic diversity of parasitic dinoflagellates in the genus amoebophrya and its relationship to parasite biology and biogeography. J Eukaryot Microbiol 55, 1–8 (2008).

36. B. E. Suzek et al., UniRef clusters: a comprehensive and scalable alternative for improving sequence similarity searches. Bioinformatics 31, 926–932 (2015).

37. N. A. O’Leary et al., Reference sequence (RefSeq) database at NCBI: current status, taxonomic expansion, and functional annotation. Nucleic Acids Res 44, D733–745 (2016).

38. B. S. Leander, P. J. Keeling, Morphostasis in alveolate evolution. Trends Ecol Evol 18, 395–402 (2003).

39. Y. H. Woo et al., Chromerid genomes reveal the evolutionary path from photosynthetic algae to obligate intracellular parasites. Elife 4, e06974 (2015).

40. S. Farhat et al., Rapid protein evolution, organellar reductions, and invasive intronic elements in the marine aerobic parasite dinoflagellate Amoebophrya spp. BMC Biol 19, 1 (2021).

41. S. Harðardóttir, et al., Millennial-scale variations in Arctic sea ice are recorded in sedimentary ancient DNA of the microalga Polarella glacialis. Commun Earth Environ 5 (2024).

42. D. R. Bogema et al., Draft genomes of Perkinsus olseni and Perkinsus chesapeaki reveal polyploidy and regional differences in heterozygosity. Genomics 113, 677–688 (2021).

43. E. Schulz et al., Crystal structure of an intramolecular chaperone mediating triple-beta-helix folding. Nat Struct Mol Biol 17, 210–215 (2010).

44. Z. Li, Y. Park, E. Marcotte, A Bacteriophage tailspike domain promotes self-cleavage of a human membrane-bound transcription factor, the myelin regulatory factor MYRF. PLoS Biol 11, e1001624 (2013).

45. D. Kim et al., Homo-trimerization is essential for the transcription factor function of Myrf for oligodendrocyte differentiation. Nucleic Acids Res 45, 5112–5125 (2017).

46. P. Haynes, Phosphoglycosylation: a new structural class of glycosylation? Glycobiology 8, 1–5 (1998).

47. G. Gustafson, L. Milner, Occurrence of N-acetylglucosamine-1-phosphate in proteinase I from Dictyostelium discoideum. J Biol Chem 255, 7208–7210 (1980).

48. N. Chandel, Carbohydrate Metabolism. Cold Spring Harb Perspect Biol 13 (2021).

49. H. Staudinger, About cellulose, starch and glycogen. Naturwissenschaften 25, 673–681 (1937).

50. P. Roach, Glycogen and its metabolism. Curr Mol Med 2, 101–120 (2002).

51. A. Smith, S. Zeeman, Starch: A Flexible, Adaptable Carbon Store Coupled to Plant Growth. Annu Rev Plant Biol 71, 217–245 (2020).

52. A. Heredia, A. Jimenez, R. Guillen, Composition of plant cell walls. Z Lebensm Unters Forsch 200, 24–31 (1995).

53. J. Archibald, Cryptomonads. Curr Biol 30, R1114–R1116 (2020).

54. Q. zhang, et al., Absolute hand determination of glycofibrils from natural sources in cryo-EM. BioRxiv (2025).

55. Q. zhang et al., Chirality determination of glycofibrils from natural sources in cryo-EM. BioRxiv (2025).

56. A. Punjani, J. Rubinstein, D. Fleet, M. Brubaker, cryoSPARC: algorithms for rapid unsupervised cryo-EM structure determination. Nat Methods 14, 290–296 (2017).

57. P. Rosenthal, R. Henderson, Optimal determination of particle orientation, absolute hand, and contrast loss in single-particle electron cryomicroscopy. J Mol Biol 333, 721–745 (2003).

58. P. Emsley, K. Cowtan, Coot: model-building tools for molecular graphics. Acta crystallographica section D: biological crystallography 60, 2126–2132 (2004).

59. P. Adams et al., PHENIX: a comprehensive Python-based system for macromolecular structure solution. Acta Crystallographica Section D: Biological Crystallography 66, 213–221 (2010).

60. I. Davis, L. Murray, J. Richardson, D. Richardson, MOLPROBITY: structure validation and all-atom contact analysis for nucleic acids and their complexes. Nucleic Acids Res 32, W615–619 (2004).

61. E. Pettersen et al., UCSF ChimeraX: Structure visualization for researchers, educators, and developers. Protein Sci 30, 70–82 (2021).

