## Supplementary material for "CryoSeek identification of glycofibrils with diverse compositions and structural assemblies": SI

**This PDF file includes:**

Figures S1-S6

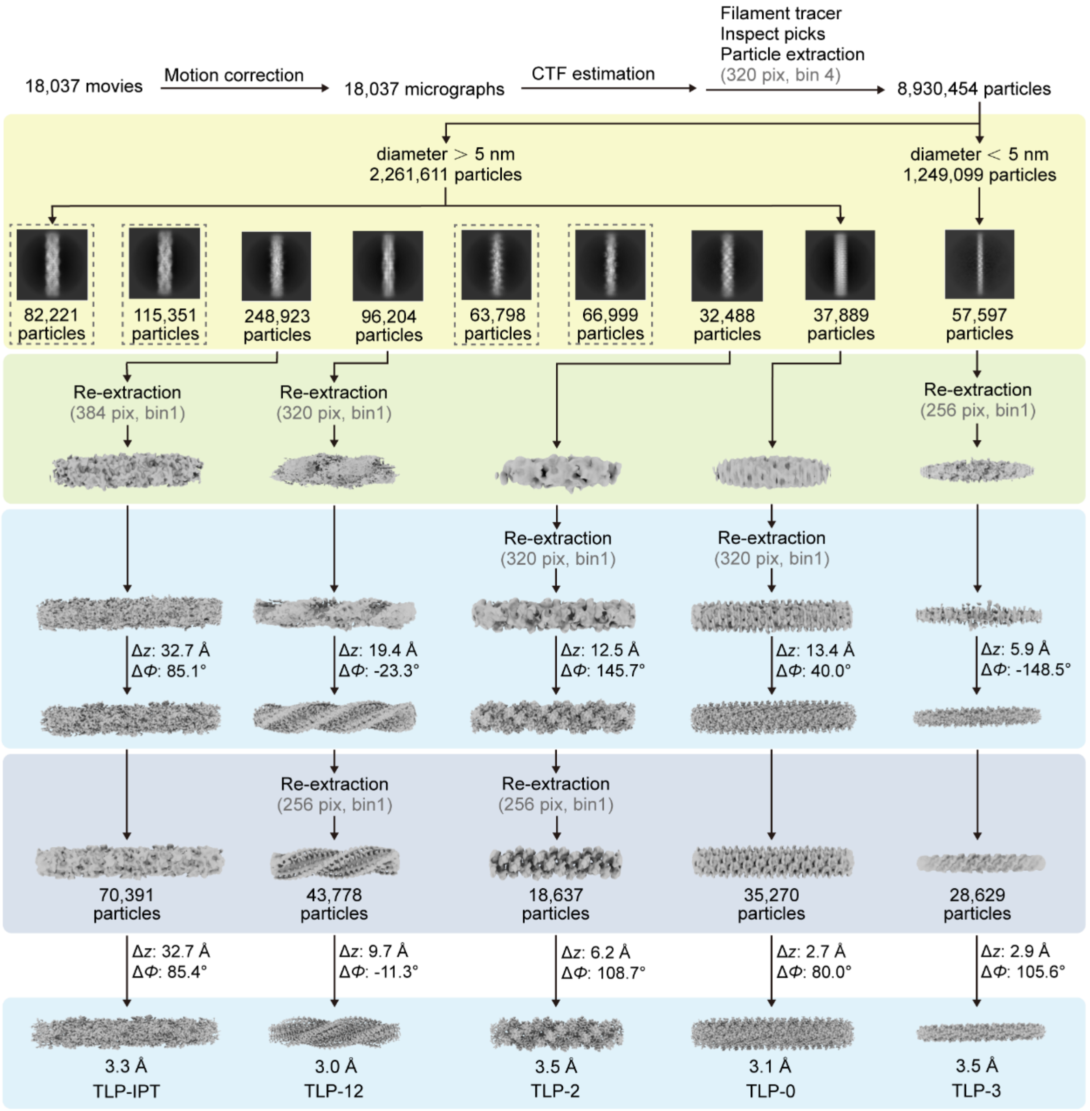


**Figure S1 | Flowchart for cryo-EM data processing of TLP fibrils.**

Details can be found in Materials and Methods. Different background colors denote distinct processing procedures: light yellow, 2D classification; light green, ab initio reconstruction; sky blue, helical refinement; and light purple, heterogeneous refinement.

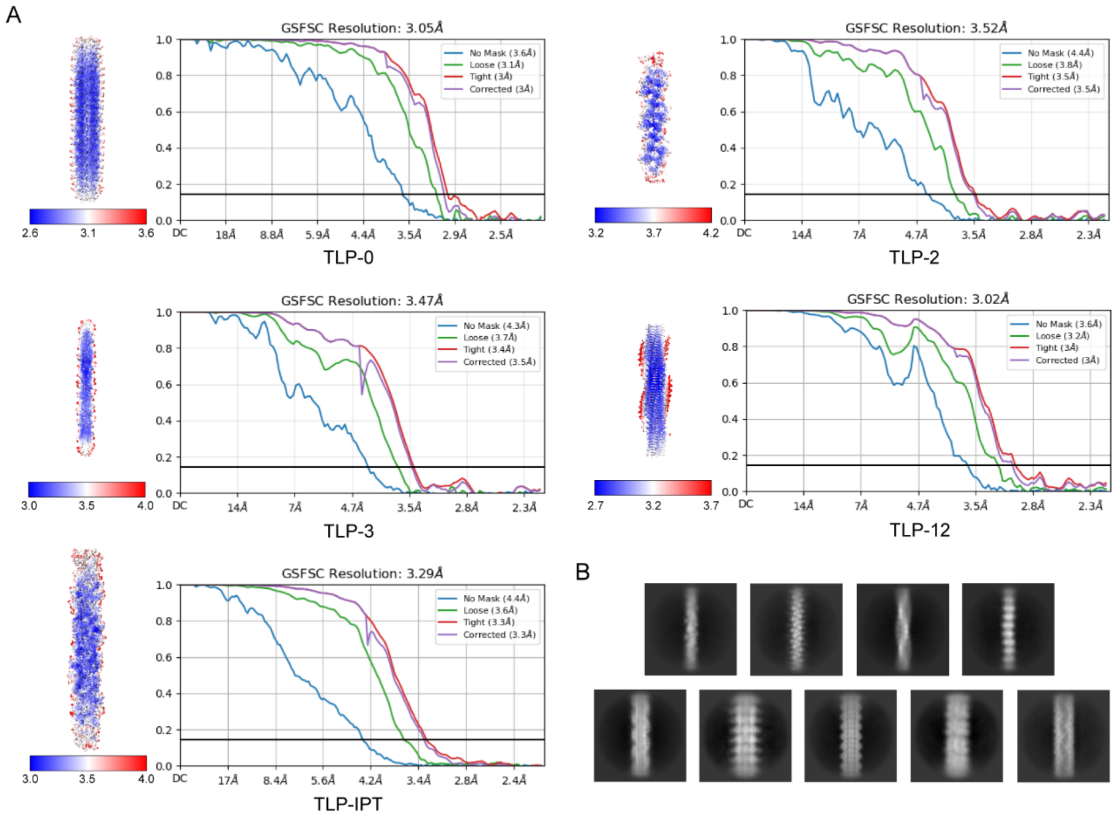


**Figure S2 | Cryo-EM analysis of TLP fibrils.**

(*A*) Local resolution distribution and gold-standard Fourier shell correlation (FSC) curves for the 3D reconstructions of TLP-0, TLP-2, TLP-3, TLP-12 and TLP-IPT. (*B*) 2D class averages of other fibrils that exhibit distinct features but currently do not support high-resolution 3D reconstructions.

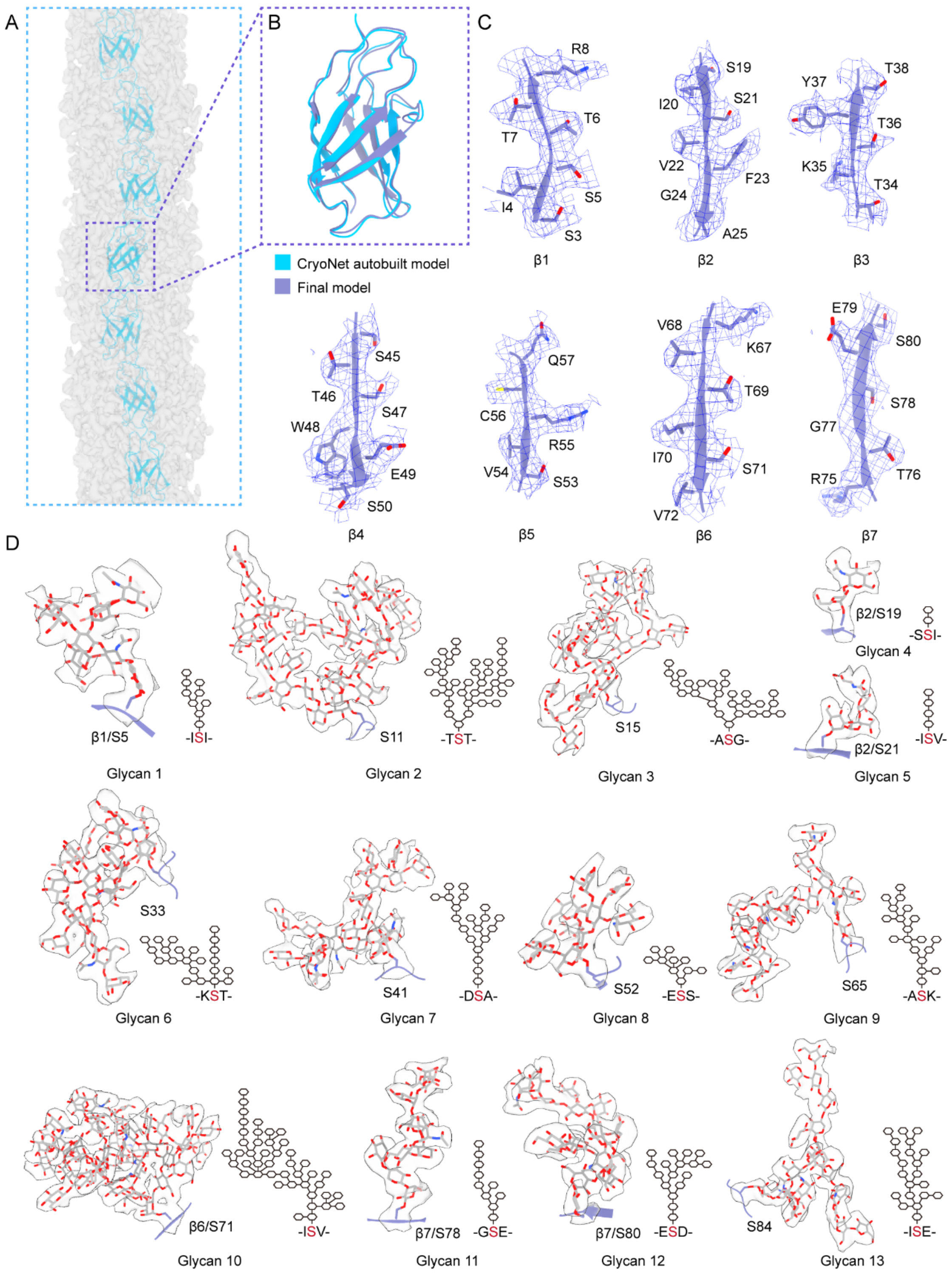


**Figure S3 | Structural model building of TLP-IPT.**

(*A*) CryoNet auto-built atomic model of the protein core of TLP-IPT. The surrounding unmodeled densities correspond to glycans. (*B*) Superimposition of the CryoNet auto-built model with the manually refined final model. The two models adopt nearly identical overall conformations. (*C*) Representative densities of β strands in a single repeat. (*D*) Tentative assignment of 13 glycan chains within one repeat. Alongside the representative densities, scheme illustrations of the glycans are presented. The EM maps are contoured at 6 σ.

A

| Species | GenBank ID | Total score | E-value | Identity (%) | Length (aa) |
|---|---|---|---|---|---|
| *Guillardia theta* CCMP2712 | XP_005822609.1 | 103 | 2e-07 | 39.47 | 1,056 |
| *Baffinella frigidus* | KAJ1470089.1 | 366 | 3e-08 | 36.05 | 1,063 |
| | KAJ1465477.1 | 207 | 2e-06 | 33.77 | 882 |
| | KAJ1463629.1 | 54.3 | 1e-06 | 44.44 | 175 |
| | KAJ1471746.1 | 282 | 1e-09 | 40.54 | 980 |
| | KAJ1467355.1 | 108 | 2e-07 | 43.24 | 307 |
| | KAJ1467356.1 | 157 | 6e-06 | 36.36 | 663 |
| | KAJ1468138.1 | 105 | 2e-06 | 36.47 | 629 |
| bacterium | MFN9897994.1 | 64.3 | 6e-11 | 37.66 | 122 |
| *Promethearchaeia archaeon* | MGB1603001.1 | 57.0 | 5e-07 | 32.47 | 377 |

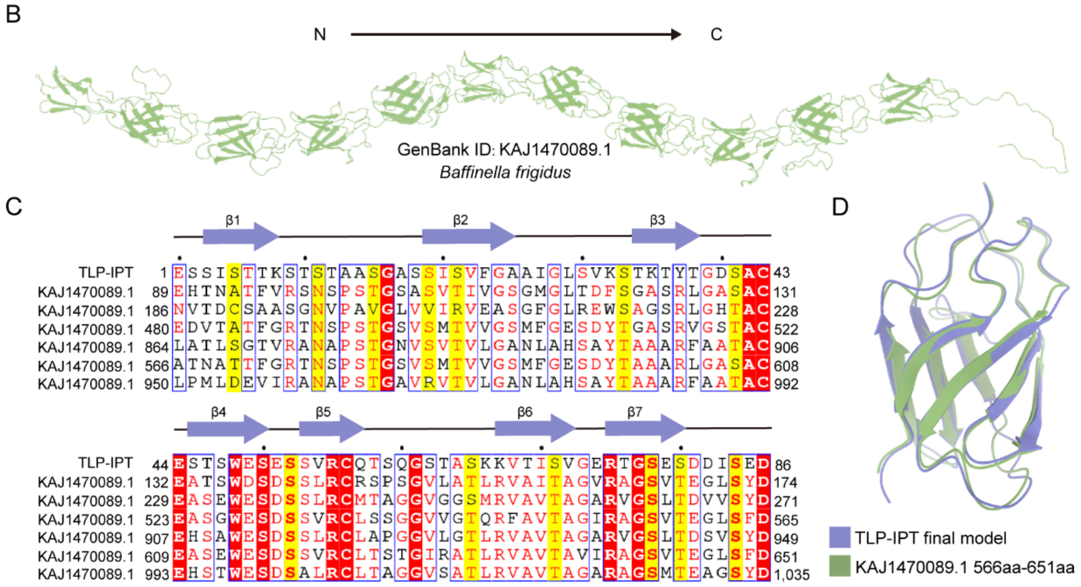


**Figure S4 | Sequence-based analysis identifies potential candidates for TLP-IPT.** (*A*) Sequences retrieved from NR database using BLAST. The refined sequence of TLP-IPT was subjected to BLAST search, a set of candidates from *Baffinella frigidus* exhibit high sequence identity to TLP-IPT, with KAJ1470089.1 displaying the highest BLAST score. (*B*) AlphaFold3-predicted structure of KAJ1470089.1, composed of consecutive similar domains, highly resembles the protein core of TLP-IPT. (*C*) Sequence alignment of one repeat in TLP-IPT with several domains of KAJ1470089.1. In addition to sequence identity, the glycosylation sites are also conserved among these repeats. (*D*) Superimposition of structures of one TLP-IPT repeat with KAJ1470089.1 (residues 566-651) reveals a nearly identical conformation.

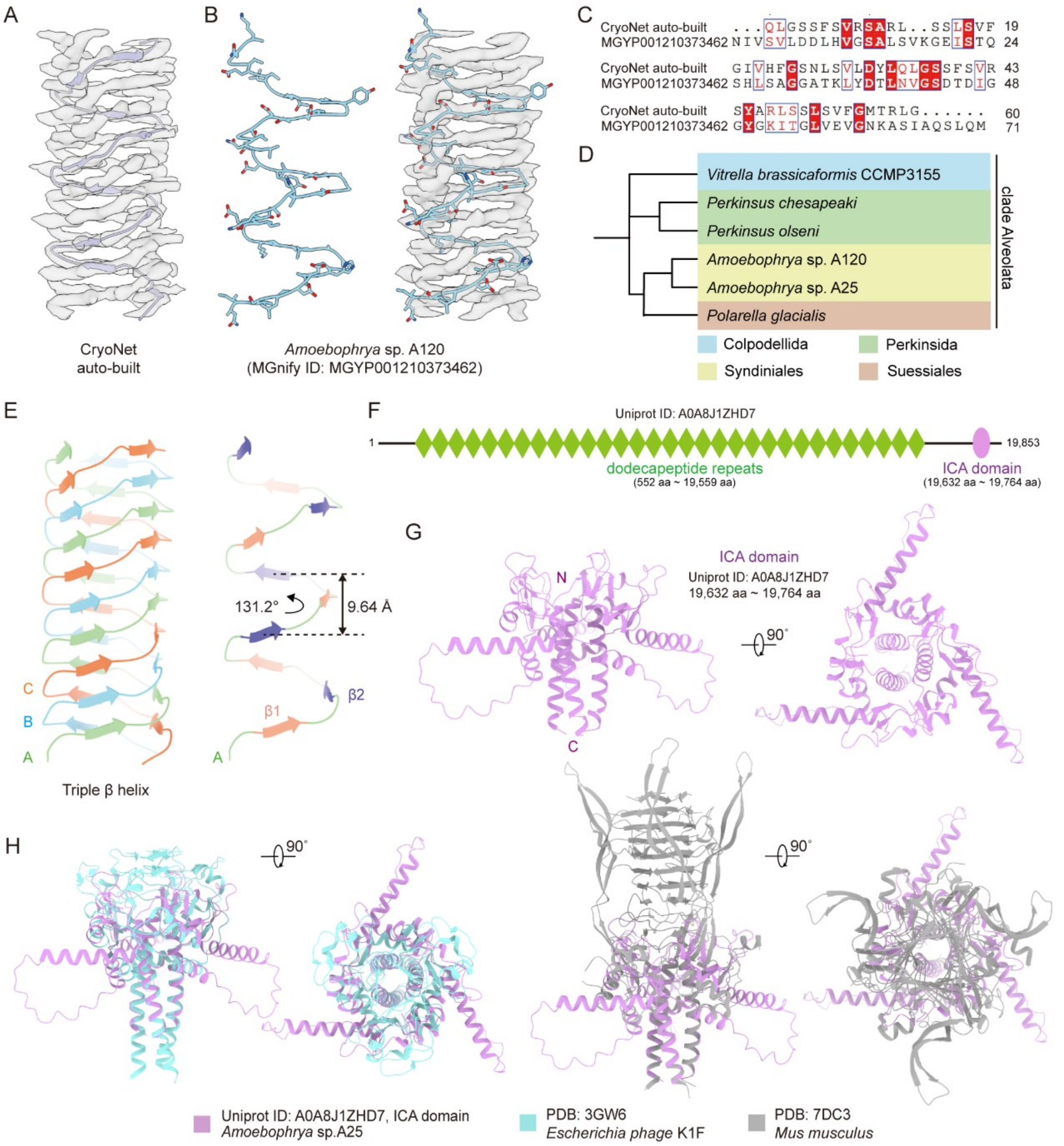


**Figure S5 | Attempt to identify the potential origins and functions of TLP-12.**

(*A*) Automatic model building for a single chain of TLP-12 using CryoNet. All glycan densities in TLP-12 were erased from the EM map to enable recognition of the amino acid residues. (*B*) Predicted model with similar fold found in ESM30 [14]. The model built in CryoNet was submitted to ESM30 for search. The sequence, with MGnify ID of MGY001210373462, has an *Amoebophrya* sp. A120 origin. The predicted atomic model fits well with the glycan-erased map. (*C*) Sequence alignment of the auto-built TLP-12 and MGYP001210373462. (*D*) Phylogenetic analysis of sequences found from NCBI NR database search via FIMO. All retrieved sequences are classified to clade Alveolata. (*E*) TLP-12 is a trimer of dodecapeptide repeats. *Left*:

The two short β strands in each dodecapeptide repeat of the three monomers, designated A, B, and C, align alternately to form three β-sheet ribbons that spiral up to constitute the central stem for TLP-12. *Right*: Two perpendicular views of a chain A segment, with β1 and β2 in each repeat being colored orange and purple, respectively. For a single chain, the helical rise and twist between two adjacent repeats are 9.64 Å and 131.2°, respectively. (*F*) Domain prediction of A0A8J1ZHD7 using InterPro reveals numerous concentrated dodecapeptide repeats in the middle region and an ICA domain at the C terminus. (*G*) AlphaFold3-predicted structure of the ICA domain from A0A8J1ZHD7. (*H*) Structural comparison of the ICA domain from A0A8J1ZHD7 with those from *Escherichia phage* K1F (PDB: 3GW6), or that from *Mus musculus* (PDB: 7DC3) reveals similar conformations.

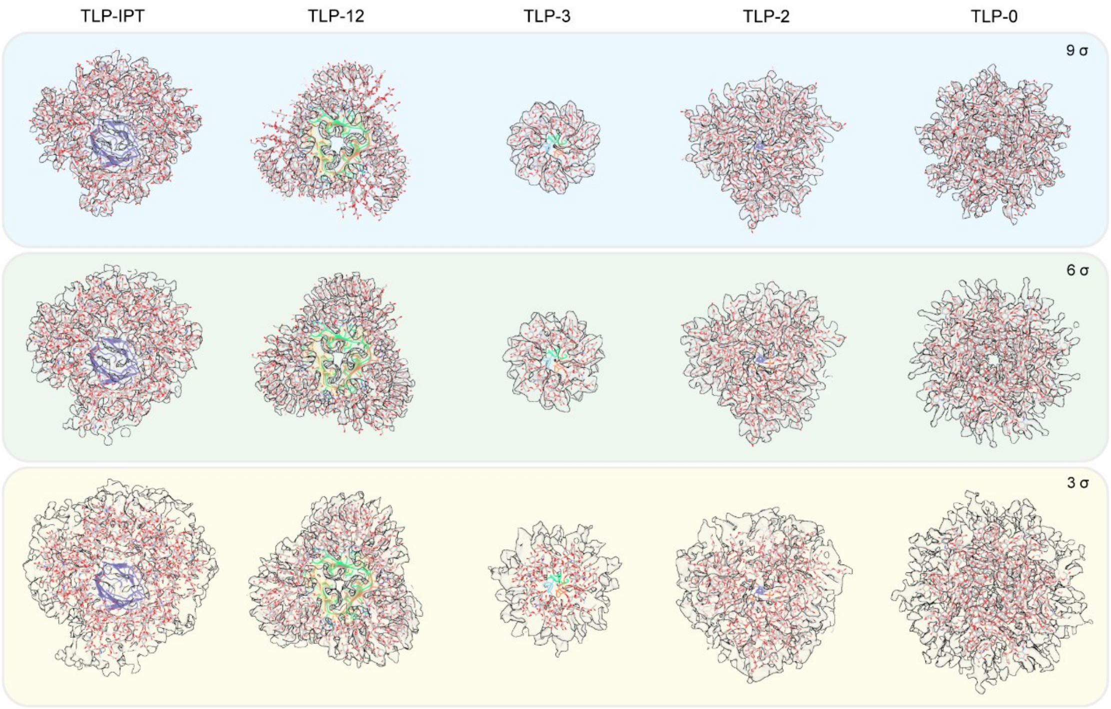


**Figure S6 | Sugar composition in the TLP glycofibrils exceeds that presented in the models.**

The resolution gradually decreases from the central axis of the fibril toward the periphery. The maps, shown in the cross-section view, are contoured at 9σ, 6σ, and 3σ from top to bottom. At lower contour thresholds, there are still many unmodeled densities.
